# Transcriptome-wide association study in UK Biobank Europeans identifies associations with blood cell traits

**DOI:** 10.1101/2021.08.03.453690

**Authors:** Bryce Rowland, Sanan Venkatesh, Manuel Tardaguila, Jia Wen, Jonathan D Rosen, Amanda L Tapia, Quan Sun, Mariaelisa Graff, Dragana Vuckovic, Guillaume Lettre, Vijay G. Sankaran, Alexander P. Reiner, Nicole Soranzo, Jennifer E. Huffman, Georgios Voloudakis, Panos Roussos, Laura Raffield, Yun Li

## Abstract

Previous genome-wide association studies (GWAS) of hematological traits have identified over 10,000 distinct trait-specific risk loci, but the underlying causal mechanisms at these loci remain incompletely characterized. We performed a transcriptome-wide association study (TWAS) of 29 hematological traits in 399,835 UK Biobank (UKB) participants of European ancestry using gene expression prediction models trained from whole blood RNA-seq data in 922 individuals. We discovered 557 TWAS signals associated with hematological traits distinct from previously discovered GWAS variants, including 10 completely novel gene-trait pairs corresponding to 9 unique genes. Among the 557 associations, 301 were available for replication in a cohort of 141,286 participants of European ancestry from the Million Veteran Program (MVP). Of these 301 associations, 199 replicated at a nominal threshold (*α* = 0.05) and 108 replicated at a strict Bonferroni adjusted threshold (*α* = 0.05/301). Using our TWAS results, we systematically assigned 4,261 out of 16,900 previously identified hematological trait GWAS variants to putative target genes. Compared to *coloc*, our TWAS results show reduced specificity and increased sensitivity to assign variants to target genes.

## Introduction

Blood cells facilitate key physiological processes in human health such as immunity, oxygen transport, and clotting. Blood cell traits have been associated with risk for complex diseases, including asthma, autoimmune conditions, and cardiovascular disease. Genome-wide association studies (GWAS) in both large European and trans-ethnic cohorts have identified thousands of loci associated with hematological traits including red blood cell, white blood cell, and platelet indices.^1–3^

While variant-level analyses provide general insights into the genetic architecture of blood cell traits, functional mechanisms for these mostly non-coding signals remain elusive. Transcriptome-wide association studies (TWAS) have been successful in identifying new genetic loci and prioritizing potential causal genes at known loci for many complex traits ^4–7^. TWAS associates phenotypes of interest with gene expressions predicted from genotype-based prediction models built in a reference eQTL dataset. TWAS results can lead to an increased understanding of the functional mechanisms underlying previously observed variant-trait associations by positing relationships between genetic variants, effector gene(s), and phenotypes. Additionally, TWAS has increased statistical power compared to single variant association tests by aggregating multiple modest strength single variant signals into a combined test^8^. Here, we conducted a large TWAS of 29 hematological traits by studying 399,835 participants of European ancestry from the UK Biobank (Figure 1)^9^. First, we trained gene expression prediction models using a reference dataset of 922 participants of European ancestry from the Depression Genes and Networks (DGN) cohort with both genotype and RNA-seq data from whole blood^10^. Second, we applied the gene expression prediction models trained in DGN to our discovery UK Biobank participants (n=399,835) to obtain predicted gene expression levels and performed association testing between predicted gene expression values and blood cell phenotypes. Third, we attempted to replicate associations identified in UK Biobank in 141,286 European ancestry participants from the Million Veteran Program (MVP) study.^11^ Finally, we performed follow-up analyses including conditional association tests on known GWAS variants, fine-mapping of TWAS loci, and TWAS-based gene assignment for GWAS variants. We demonstrate advantages of TWAS over single-variant analyses by comparing to a recent large GWAS of hematological traits in UK Biobank Europeans^3^.

**Figure 1.**
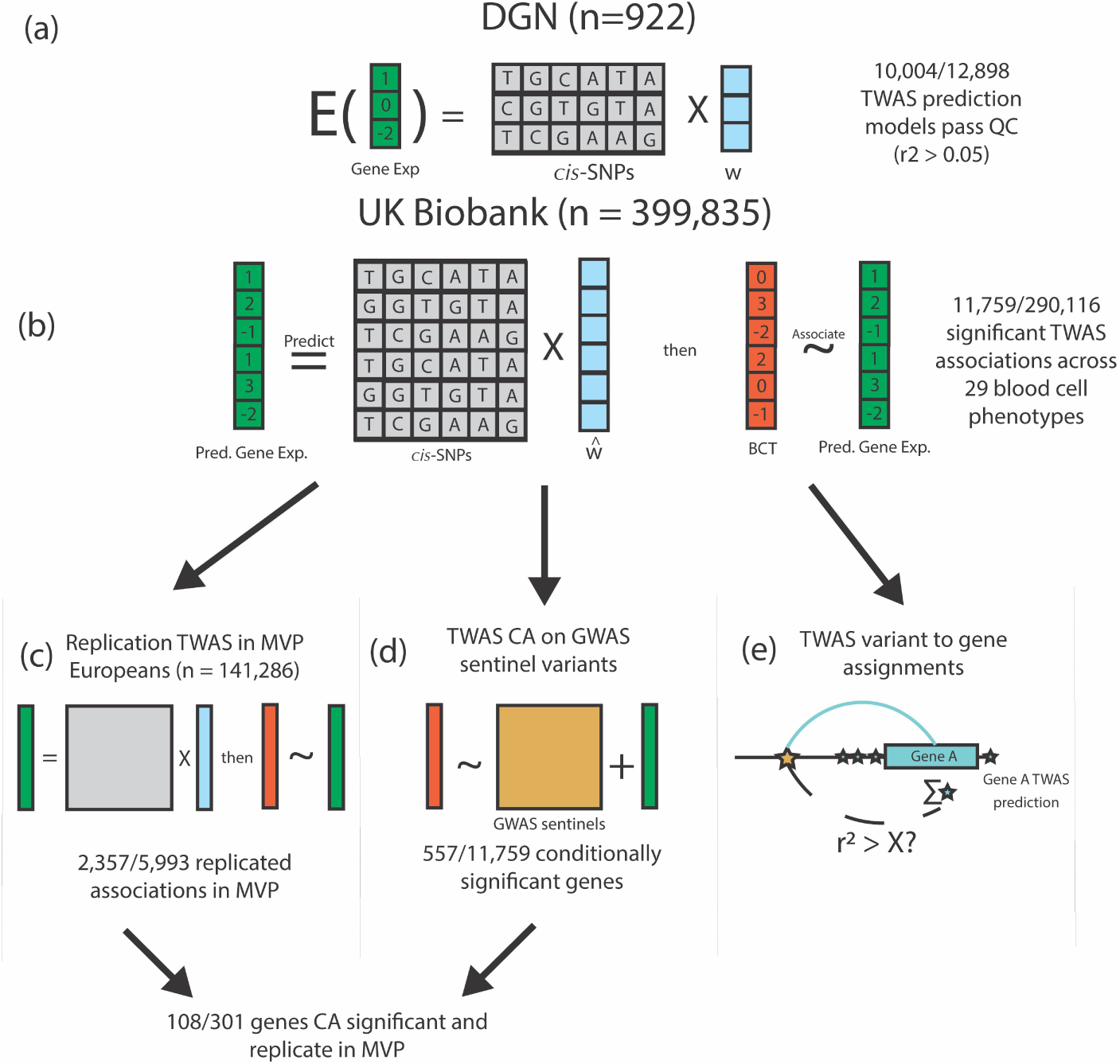
UK Biobank TWAS of Blood Cell Traits Overview. (a) We trained gene expression prediction models using whole blood gene expression data from 922 Depression Genes and Networks (DGN) European ancestry participants by fitting an elastic net model on the *cis*-SNPs (+/-1Mb) for each gene. Models with r^2^ > 0.05 are considered sufficiently predicted, and are subject to association testing in UKB. *w* represents the TWAS weights in the prediction model. (b) Using our DGN-trained models, we predicted gene expression in 399,835 UKB participants of European ancestry and performed association testing with 29 hematological traits. 11,759 gene-trait associations were significant at the Bonferroni adjusted threshold (out of 290,116 tested). (c) TWAS results from UKB were replicated in 141,286 MVP participants of European ancestry for 15 hematological traits available in MVP. (d) We further conditioned our TWAS significant associations in UKB on GWAS signals reported from Vuckovic *et al.* to determine which TWAS signals were driven by previously reported GWAS variants (TWAS CA for TWAS conditional analysis). (e) We used the TWAS associations and prediction models (blue stars) to assign GWAS signals from Vuckovic *et al.* (gold star) to plausible target genes (see Figure 2), assessing correlation between each GWAS variant and predicted gene expression of each TWAS significant gene.

As previously mentioned, TWAS results can shed light on the functional mechanisms underlying variant-trait associations by linking variants to target genes. Designing appropriate functional experiments to interrogate biological mechanisms or to identify potential drug targets necessitates accurately assigning GWAS variants to target genes. Often, variants are linked to target genes using distance based approaches, which can lead to inaccurate assignments (“Nearest Gene” in Figure 2).^12, 13^ Colocalization based methods (“eQTL Colocalization” in Figure 2) evaluate the evidence that a GWAS variant coincides with an eQTL signal for a gene in a relevant cell type and if these signals are likely driven by the same biological process or the same set of variants. While useful, colocalization methods may be underpowered in situations where there are multiple variants which are associated both with a complex trait in GWAS and linked to the same target gene but with low or moderate effect size.

**Figure 2.**
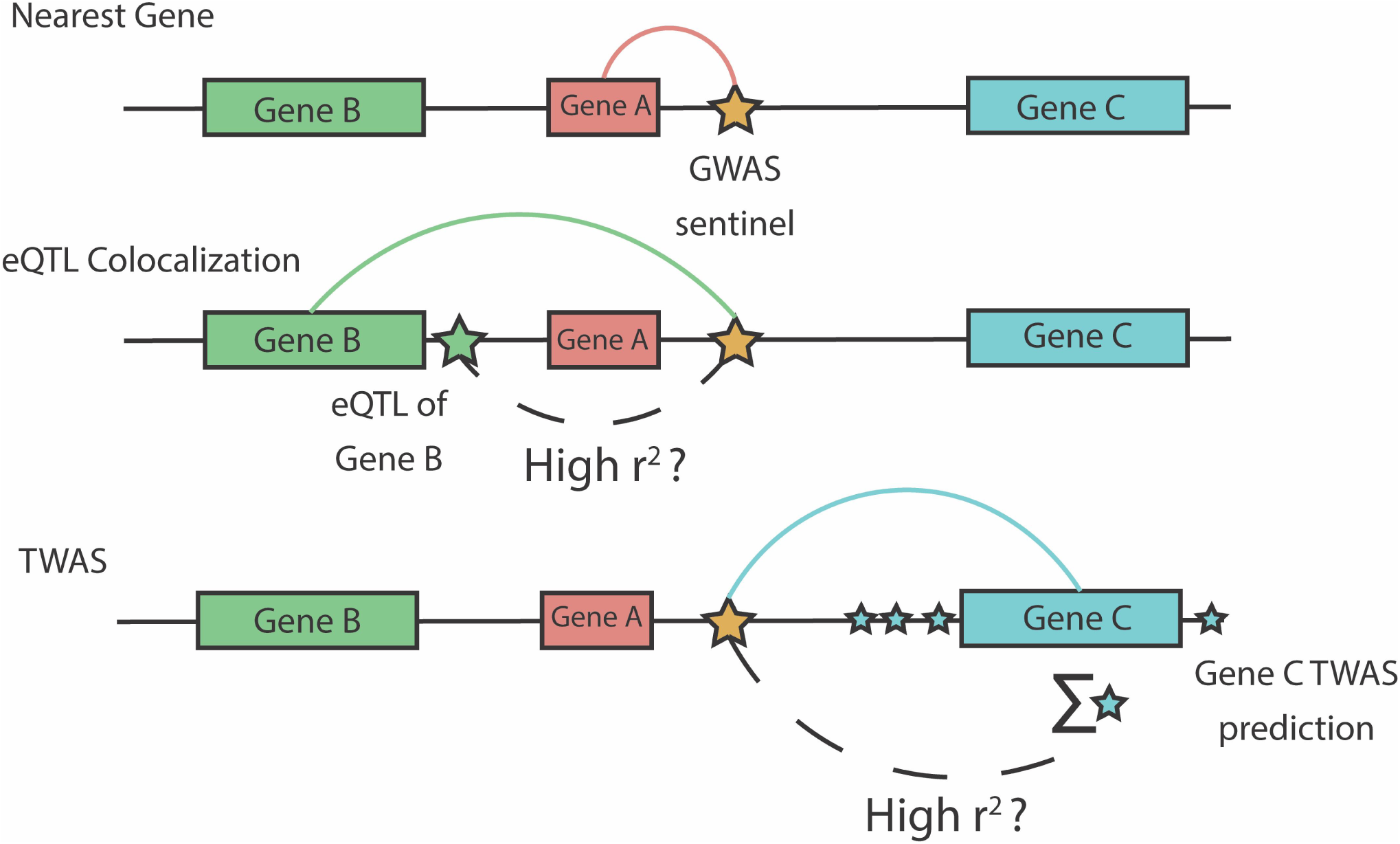
TWAS variant-to-gene Approach. Comparison of our TWAS-based approach to variant-to-gene assignment with two commonly used approaches: distance-based and colocalization based assignments. We consider the problem of assigning a GWAS variant (gold star) in a non-coding region to a target gene. The nearest-gene approach assigns the variant to the closest gene at the locus (Gene A), but ignores epigenomic evidence at the locus. Colocalization based approaches assign the variant to a target gene based on evidence that the GWAS signal is not distinct from an eQTL signal for a target gene (green star, Gene B). Our TWAS based approach assesses the correlation between the GWAS variant and TWAS predicted gene expression which aggregates smaller effect *cis*-eQTLs for a gene (blue stars, Gene C). For presentation brevity, we use “high r^2^” but the threshold to define high correlation can be lenient.

We leveraged our TWAS results to assign GWAS variants to target genes. For each GWAS variant-trait association, our TWAS based approach identified a set of potential target genes associated with the same phenotype utilizing TWAS association results and gene expression prediction models (“TWAS” in Figure 2). We then used individual level genotypes in our testing cohort (UK Biobank) to identify variant-gene pairs where the variant genotype and the predicted gene expression values are correlated (r2 > 0.2). By effectively aggregating multiple smaller effect eQTLs for a gene, we hypothesized that our TWAS based approach would be better powered than *coloc* to identify target gene(s) for a GWAS variant. We systematically assigned the 16,900 conditionally distinct variant-trait associations identified by Vuckovic *et al.* to target genes and compared our TWAS-based assignments to those from *coloc*, a commonly used eQTL colocalization method.

## Methods

### Included Cohorts

#### Depression Genes and Networks (DGN)

The DGN study was designed to collect samples of individuals with and without major depressive disorder, ages 21-60, from a survey research panel broadly representative of the United States population.^10^. Genotyping and RNA-sequencing procedures have been described previously.^10^ For 922 European ancestry participants from the DGN study, we obtained both genotype data imputed to the TOPMed Freeze 8 reference panel and RNA-seq data.^14, 15^ For training gene expression prediction models, we included bi-allelic variants that are common and well-imputed (MAF > 0.05, Rsq > 0.8) in both DGN and in the UK Biobank. In all, 5,652,397 variants were included, here forward referred to as QC variants. DGN whole-blood RNA-seq data was obtained for 922 European ancestry participants.^10^ As described previously, quantified gene expression values were normalized using the hidden covariates with prior (HCP) method^16^, correcting for technical and biological factors.^10^

#### UK Biobank (UKB) Europeans

UKB recruited 500,000 people aged between 40-69 years in 2006-2010, establishing a prospective biobank study to understand risk factors for common diseases such as cancer, heart disease, stroke, diabetes, and dementia.^9^ Participants are being followed-up through health records from the UK National Health Service. UKB has genotype data, imputed with UK10K as reference, on all enrolled participants, as well as extensive baseline questionnaires and physical measures and stored blood and urine samples. Hematological traits were assayed as previously described.^1^ Genotyping on custom Axiom arrays, subsequent quality control, and imputation has been previously described.^9^

For our TWAS, we analyzed UKB participants of European ancestry to match the genetic ancestry of DGN participants used for model training. Participants were included in our analysis if identified as European through a combination of self-reported ancestry and *k*-means clustering of genetic principal components (PCs) in order to minimize genomic inflation due to population stratification, and for consistency with previously published blood cell trait GWAS in UKB.^3^ First, we calculated PCs and their loadings for all 488,377 genotyped UKB participants using LD pruned variants (pairwise r^2^ < 0.1) with MAF ≥0.01 and missing rate ≤0.015 in the UK Biobank data set that overlapped with the participants in the 1000G Phase 3 v5 (1KG) reference panel.^9^ Reference ancestries used included 504 European (EUR), 347 American (AMR), 661 African (AFR), 504 East Asian (EAS) and 489 South Asian (SAS) samples (overall 2504). We projected the 1KG reference panel dataset on the calculated PC loadings from UK Biobank. We then used *k*-means clustering with 4 dimensions, defined by the first 4 PCs, to identify individuals that clustered with the majority of 1KG reference panels in each ancestry. We used self-reported ancestry/ethnicity, in some circumstances, to adjust these groups. UKB participants defined as European ancestry include those that cluster with the most 1KG EUR by *k*-means clustering. We adjusted this group by removing those that self-reported as Indian, Pakistani, Bangladeshi, any other Asian background, Black or Black British, Caribbean, African, Any other Black background, or Chinese (n=32). Additionally, we removed any individuals with self-reported mixed ancestry (n=402). A total of 451,305 remained in the European ancestry group. Participants were also excluded based on factors likely to cause major perturbations in hematological indices including positive pregnancy status, drug treatments, cancer self-report, ICD9 and ICD10 disease codes (see Supplemental Text), and surgical procedures. Participants were included only if they had complete data for all covariates and phenotypes. In total, 399,835 samples were included in the analysis.

As aforementioned, we included only bi-allelic and well-imputed common variants (Rsq > 0.8, MAF > 0.05) in UKB. All 29 blood cell phenotypes were adjusted for age, age^2^, top 10 genotype PCs, center, genotyping array, and sex. For white blood cell traits, phenotypes were log10(x + 1) transformed before regression. Residuals from these regression models were inverse normal transformed and serve as phenotypes.

#### Million Veteran Program (MVP) Europeans

The MVP is an observational cohort study and mega-biobank in the Department of Veteran Affairs healthcare system which began enrollment in 2011. As of Release 3, 318,725 individuals of European ancestry (as defined by HARE^17^) have available electronic health records (EHR), survey, and genotype data. After quality control largely following the guidelines established in Marees et al 2018, 308,778 individuals of European ancestry remained.^18^ Only a subset of 15 hematological traits out of the 29 analyzed in UKB were available for replication in MVP. For our replication study, participants were limited to those with available data among these 15 traits (n = 141,286). Phenotypes were adjusted for covariates following the same procedure as in UKB.

##### Training of Gene Expression Prediction Models

We trained gene expression prediction models using an elastic net pipeline following the well-established PrediXcan methodology.^8^ Our decision to use an in-house pipeline rather than the publicly available weights from PrediXcan was two-fold. First, we performed TOPMed freeze 8 based imputation, enhancing genome coverage and imputation quality compared to the 1000G Phase 1 v3 reference panel underlying the PrediXcan weights, the 1000G Phase 1 v3 ShapeIt2 (no singletons) panel. Second, by training our own prediction models, we ensured that every variant present in the prediction models was available in our UKB dataset.

For each gene, we included variants within a 1Mb window of the gene start and end positions and excluded variants in high pairwise LD (r^2^ > 0.9) with other variants in the window. We tuned the elastic net penalty parameters using 5 fold cross validation with the ‘glmnet’ function in R. We obtained 12,898 elastic net models where more than one variant was included in the prediction model. Models with a single variant were excluded from our TWAS since there is no difference between the TWAS approach and single variant GWAS in this setting. We further excluded models with model R2 <= 0.05, leading to 10,004 models for subsequent analysis (Figure S1).

##### Association Testing with REGENIE

Using the 10,004 models trained in DGN, we predicted gene expression values in UKB European ancestry participants. We then performed association testing between predicted gene expression and covariate-adjusted blood cell phenotypes with REGENIE.^19^ We used an LD (linkage disequilibrium) pruned (plink --indep-pairwise 50 5 0.1) set of 174,957 variants with MAF > 0.01 in the genotype data available for UKB Europeans to fit the REGENIE null model accounting for cryptic relatedness. We analyze all 29 phenotypes simultaneously using the grouping option available in REGENIE and set the number of blocks to 1,000.

To control Type I error at *α* = 0.05, we considered a TWAS association significant if p < 0.05/(10,004 * 29) = 1.72*10^-7^. Note that this Bonferroni adjusted significance threshold is rather quite conservative due to correlations among the blood cell phenotypes and among predicted expressions of genes. Results from this TWAS association analysis are referred throughout the manuscript as the marginal TWAS results. After the marginal analysis, we partitioned the results into TWAS loci for each trait by beginning with the most significant TWAS gene not assigned to a TWAS locus, assigning all genes within 1Mb of the gene not yet in a locus to its locus, and then proceeded similarly for the entire genome until all significant genes were in a locus.

##### Conditional Analysis

In order to assess which marginally significant TWAS genes provide novel findings above and beyond the discoveries in GWAS of blood cell traits in Europeans, we tested the association between predicted gene expression and phenotype while conditioning on reported blood cell trait GWAS variants. This methodology has been described in a previous TWAS of blood cell traits from our group.^20^ We partitioned the distinct GWAS variants from Vuckovic *et al.* into three phenotype categories: red blood cell (RBC), white blood cell (WBC), and platelet (PLT) traits.^3^ We considered all distinct GWAS variants as determined by conditional analysis on individual level data, referred to as conditionally independent variants by Vuckovic *et al.* For a TWAS gene associated with one trait in the above categories, we conditioned on any distinct variant reported as associated with any trait within the corresponding phenotype category on the same chromosome.

##### Replication Analysis in MVP

We conducted two replication analyses in MVP Europeans to follow up on our results from the UKB TWAS: one for the marginal TWAS results and a second restricted to only conditionally significant genes. In both analyses, our DGN trained gene expression prediction models were used to impute gene expression values in MVP Europeans. Association testing was performed via boltLMM.^21^ The Bonferroni adjusted thresholds for replication were determined by the number of marginal or conditional associations in the UKB available for replication, respectively.

##### TWAS fine mapping via FINEMAP

We modified the FINEMAP software to compute credible sets of genes from our marginal TWAS results.^22^ We substituted GWAS summary statistics for our TWAS summary statistics from the marginal TWAS analysis. In place of an LD matrix, we used a gene-gene correlation matrix computed on the predicted gene expression values in UKB Europeans. We compute the FINEMAP credible sets and posterior probabilities of inclusion for all TWAS loci with at least 2 genes.

### TWAS variant-to-gene assignments

We assigned the distinct GWAS variants from Vuckovic *et al.* to putative target genes using our TWAS results.^3^ For a GWAS variant-trait association, we considered all significant TWAS genes for the matching trait in any TWAS locus within 1Mb of the variant. We assigned the variant to a gene if the TWAS gene had both a FINEMAP posterior probability of inclusion greater than 0.5, and evidence of correlation (r^2^ > 0.2) between the variant genotype and predicted gene expression. We performed our TWAS assignments on 10,239 variant trait associations across 10 hematological traits from Vuckovic *et al*.^3^ These 10 traits were chosen based on data availability for eQTLs in relevant cell types including platelets, CD4+, CD8+, CD14+, CD15+, and CD19+ cells. In their work, they performed eQTL co-localization analyses using *coloc*. For a GWAS variant, we assigned the eGene(s) corresponding to any co-localizing eQTL as the target gene.

#### Open Targets

Open Targets Genetics is an open-access integrative resource which aggregates human GWAS and functional genomics data including gene expression, protein abundance, chromatin interaction, and conformation data in order to make robust connections between GWAS loci and potentially causal genes.^23^ In order to assign potentially causal genes to a given GWAS variant, Open Targets provides a disease-agnostic variant-to-gene (V2G) score which combines a single aggregated score for each GWAS variant-gene prediction. This analysis combines four different data types: eQTL and pQTL datasets, chromatin interaction and conformation datasets, Variant Effect Predictor (VEP) scores, and distance from the canonical transcription start site for a target gene. We compare the TWAS and *coloc* variant-to-gene assignments to the sets of potentially causal genes identified by Open Targets. Performance is assessed by checking if any TWAS/*coloc* assigned gene for a given variant is either the most likely gene identified by Open Targets (OT Max) or any gene identified by Open Targets (OT Any).

#### BLUEPRINT specifically expressed genes

We also assessed the quality of the gene assignments for the TWAS and *coloc*-based methods by determining if the assigned gene is cell-type specifically expressed in gene expression data from BLUEPRINT.^24^ We group available expression data into five cell type groups: erythrocytes, megakaryocytes, macrophages and monocytes, nCD4 cells, and neutrophils. We classified genes as cell type group-specific or shared via Shannon entropy across the five cell type groups. We first exponentiated the BLUEPRINT MMSEQ expression quantifications, to be comparable to RPKM. Then, for each gene, we calculated the normalized gene expression by dividing gene expression in each cell type group by the sum across all five cell type groups. Next, we calculated Shannon entropy using the normalized gene expression values. We defined the shared genes across cell type groups as those with entropy < 0.1 and the cell type-specific genes as those with entropy > 0.5 and gene expression > 1 in the respective cell type. Biologically plausible cell type groups selected for the 29 phenotypes analyzed are detailed in Supplemental Table S1.

## Results

### Marginal TWAS Results

Using an elastic net-based pipeline, we trained gene expression prediction models using imputed genotypes and whole blood RNA-seq data from 922 European ancestry participants from the DGN cohort^10^. In total, we trained prediction models for 12,989 genes, 10,004 of which passed our quality control filter (model R2 > 0.05 and >1 variant selected in model) (Supplementary Figure 1).

We conducted a TWAS in 399,835 participants of European ancestry from the UKB for 29 blood cell phenotypes: 11 white blood cell indices, 4 platelet indices, and 14 red blood cell indices (see Supplemental Table S1). 11,759 gene-trait associations were transcriptome-wide significant at the Bonferroni adjusted threshold of 1.72 ×10^-7^. The 11,759 associations were grouped into 4,835 trait-specific TWAS loci (see **Methods**) with the most significant gene at each TWAS locus assigned as the sentinel TWAS gene, resulting in 1,792 unique sentinel genes. Among these 1,792 genes, 1,112 were sentinel genes for more than one trait (see Figure S2). Of the 4,835 TWAS loci, 2,375 (49.1%) had multiple TWAS significant genes. Additionally, the 1Mb region surrounding 9 sentinel genes did not contain any distinct GWAS variants from Vuckovic *et al*.

We generated credible sets at all TWAS loci using FINEMAP (see **Methods** for details).^22^ 8,928 out of 11,759 (76%) marginal TWAS associations were included in the FINEMAP credible sets for their trait-specific loci. The average number of genes in each FINEMAP credible set was 3.97 (SD = 2.3) and the median was 4 (see Supplemental Table S2). In 297 (6.1%) trait specific loci, the sentinel TWAS gene was not included in the credible set.

In order to replicate significant results from our marginal TWAS analysis, we predicted gene expression values in 141,286 European ancestry participants from MVP using models trained in DGN (see **Methods**).^11^ For the replication analysis, 15 out of the 29 UKB analyzed blood cell traits were available in MVP. 9,492 out of the 10,004 (94.8%) gene expression prediction models were comprised of variants that overlapped completely with variants available in MVP. Replication was thus attempted in MVP for 5,993 gene-trait pairs with fully matching phenotype and gene expression prediction model variants. Among the attempted 5,993 gene-trait pairs marginally significant in UKB (marginal in contrast to conditional on nearby GWAS variants), 4,245 (71.3%) replicated in MVP at a nominal significance threshold (*α* = 0.05) with the same direction of effect, and 2,357 (39.3%) replicated at the Bonferroni corrected threshold (*α* = 8.34 ×10^-6^) with the same direction of effect (Figure S3).

### Conditional Analyses Adjusting for Nearby GWAS Variants

We then used conditional analysis to determine which of the 11,759 gene-phenotype associations in UKB represent novel findings beyond the recently published GWAS (see **Methods** for details)^3^. Of the 11,759 marginal gene-trait associations, 557 were conditionally significant at the Bonferroni corrected threshold (*α* = 0.05/11,759 = 4.25×10^-6^, Figure 3). These 557 associations represent 395 distinct genes in 463 trait-specific loci; 276 genes were conditionally significant for one trait, and 119 for multiple traits (Figure S4).

**Figure 3.**
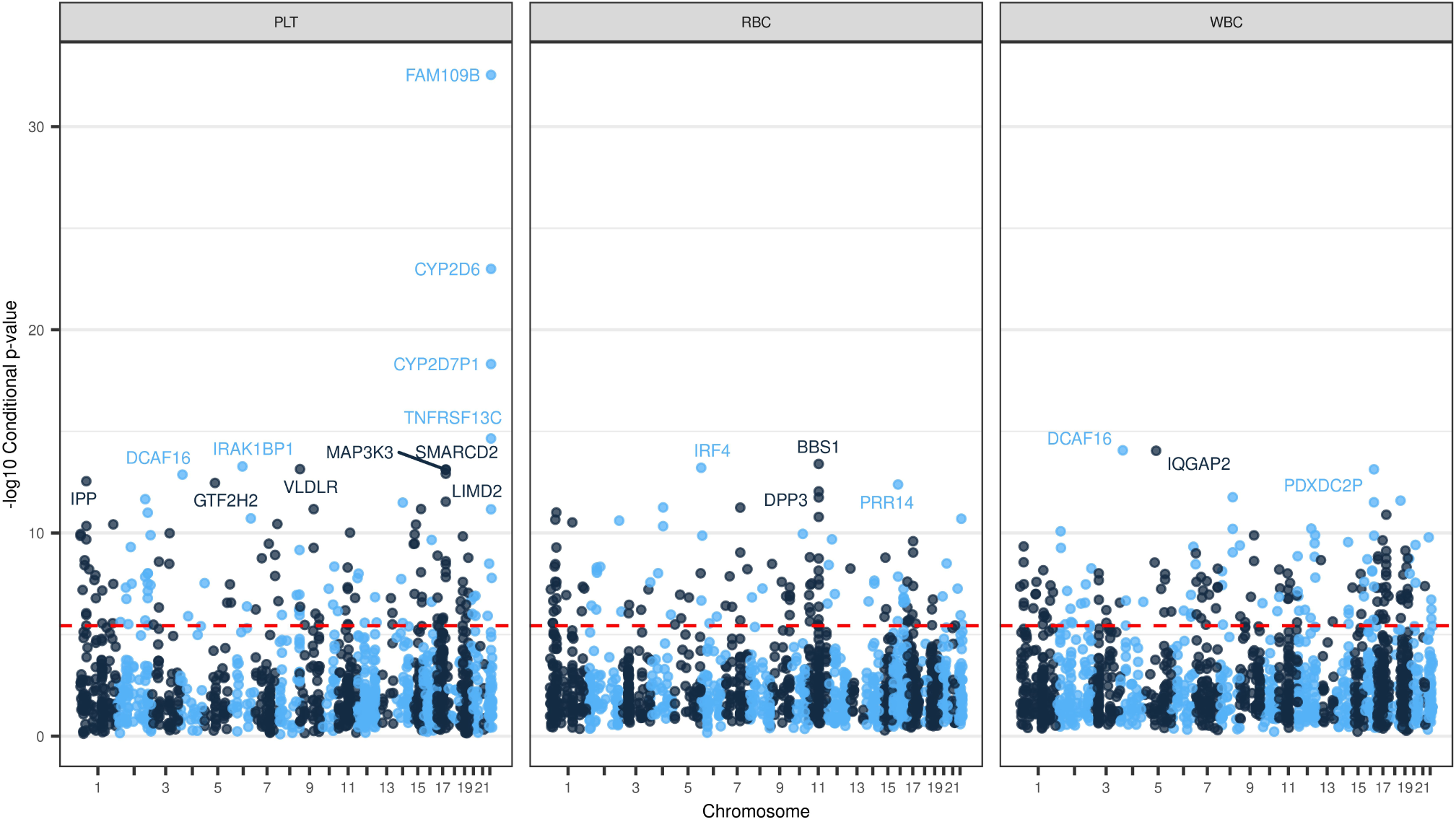
Manhattan Plot of TWAS Conditional Analysis Results. Figure 3 shows the -log10(p-value) for TWAS genes after conditioning on distinct GWAS signals from Vuckovic *et al.* for a given phenotype category. The red dashed line denotes the Bonferroni adjusted significance threshold (*α* = 4.25×10^-6^). Named genes have -log10(p-value) > 12. The conditional TWAS analysis assesses whether a TWAS signal is driven primarily from signals at previously discovered GWAS loci, which is a crucial step for our analysis of well-studied hematological traits. The maximum-log10(p-value) for each gene is plotted and stratified by phenotype category.

Of the 557 conditionally significant associations discovered from UKB, 301 had both matching phenotypes and predicted gene expression in MVP, and thus were subject for replication. 199 (66.1%) associations replicate at a nominal threshold (*α* = 0.05) with consistent direction of effect (Figure S5), and 108 associations (35.9%) replicate at a Bonferroni adjusted threshold (0.05/301 = 1.66×10^-4^) with matching direction of effect.

Additionally, 36 of the 557 conditionally significant TWAS associations had no GWAS significant (p < 5 ×10^-8^) variants for the associated trait within 1Mb of the gene. These 36 associations reflect TWAS’s increased power over single-variant GWAS analyses by aggregating multiple sub-genome-wide significant GWAS variants. For example, raftlin family member 2 (*RFTN2*) (GRCh37 chr2:198,432,948 - 198,540,769) which plays a role in TICAM-1 signaling in dendritic cells^25^, was associated with white blood cell count via marginal (p = 1.32×10^-7^) and conditional analysis (p = 2.69×10^-7^). In Vuckovic *et al.*, the sentinel variant at the locus did not achieve genome-wide significance (p = 2×10^-6^). The association between *RFTN2* and white blood cell count replicated in MVP (p = 1.9×10^-8^) with the same direction of effect.

#### Novel TWAS Loci

We discovered 10 conditionally significant gene-trait associations that have no previously identified distinct GWAS variants within ±1Mb of the locus for any blood cell trait. Among the 10 associations, 4 were unable to be assessed for replication in MVP due to phenotype unavailability or missing variants in the TWAS prediction model. 3 out of the remaining 6 associations replicated in MVP at a nominal significance threshold (*α* = 0.05) with the same direction of effect as in UKB, namely, interleukin 1 receptor associated kinase 1 binding protein 1 (*IRAK1BP1*) for mean platelet volume (beta = 0.0304, p = 3.4×10^-6^), and *SNHG5* for neutrophil count (beta = −0.0146, p = 0.0061) and white blood cell count (beta = - 0.0134, p = 0.013). The association between *IRAK1BP1* and mean platelet volume also achieved significance at the Bonferroni adjusted replication threshold.

##### *IRAK1BP1* (GRCh37 chr6:79,577,189 - 79,656,157)

The TWAS association between *IRAK1BP1* and mean platelet volume discovered by our TWAS demonstrates the utility of TWAS to discover and confirm trait-associated loci beyond GWAS and conditional analysis at the single variant level. In our TWAS, *IRAK1BP1* demonstrated evidence of association with mean platelet volume despite no conditionally distinct GWAS signals within 1Mb of the gene. The 1Mb region around *IRAK1BP1* contains several genome-wide significant variants in Vuckovic *et al.*, with lead variant GRCh37 chr6:79617522 (p=6.4×10^-13^) (Figure 4a). However, this region was grouped into a mean platelet volume locus over 8Mb away via individual-level conditional analysis (sentinel variant GRCh37 chr6:71326034_G_A). Importantly, in the Vuckovic *et al.* results, no target gene was identified for chr6:71326034_G_A via Ensembl Variant Effect Predictor (VEP)^26^ limiting the biological interpretation of the findings at the GWAS locus. Our TWAS prediction model for *IRAK1BP1* is primarily driven by variants in high LD with chr6:79617522; of the top 15 variants in terms of absolute value of the TWAS weights, 13/15 are in high LD (r2 > 0.8 in TOP-LD EUR) with chr6:79617522 (Figure 4b).

**Figure 4.**
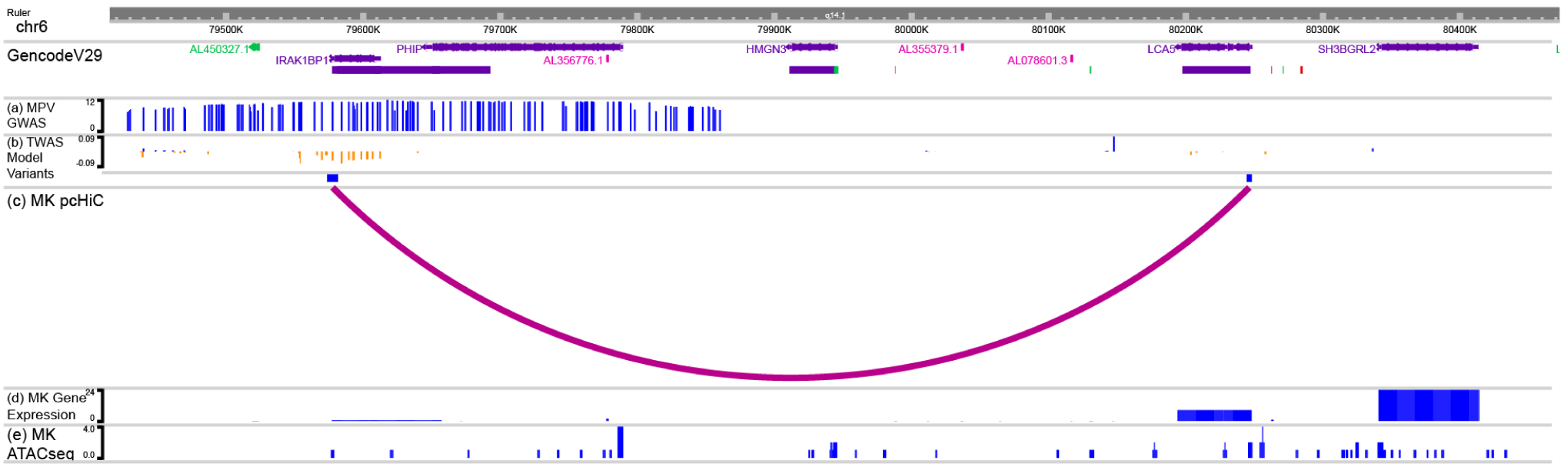
*IRAK1BP1* locus EpiGenome Browser. Figure 4 demonstrates the cell-type specific epigenetic information linking *IRAK1BP1* and mean platelet volume. Figure 4a shows that variants within *IRAK1BP1* were identified as GWAS significant variants in *Vuckovic et al., 2020*, but the signal at this locus was attenuated after conditioning on a locus 8Mb away (sentinel variant chr6:71326034_G_A). (b) Several of these variants are included in the *IRAK1BP1* TWAS prediction model. (c) Promoter-capture Hi-C data support that TWAS model variants for *IRAK1BP1* form a loop with the promoter region of *LCA5*, a mendelian disease gene for Leber congenital amaurosis. *LCA5* was not available in our DGN expression dataset. (d) *LCA5* is more strongly expressed in MK cell lines compared to *IRAK1BP1*. (e) TWAS model variants overlap with MK ATACseq peaks. This locus shows both the statistical power gains possible through TWAS analysis and the importance of full consideration of other potential target genes and complementary functional annotation resources in interpretation of a TWAS identified signal.

After conditioning on all distinct platelet-related variants on chromosome 6, including chr6:71326034_G_A, the marginal TWAS association for *IRAK1BP1* and mean platelet volume (p = 9.47×10^-12^) was not attenuated (p = 5.33×10^-14^), demonstrating that the *IRAK1BP1* TWAS signal is distinct from previously reported GWAS variants. Furthermore, the association between *IRAK1BP1* and mean platelet volume replicated in MVP Europeans at the Bonferroni adjusted threshold (p = 3.4×10^-6^). Thus, with TWAS, we combined several trait-associated variants at the *IRAK1BP1* locus into a stronger signal which demonstrated statistical independence from the previously reported chr6:71326034_G_A signal and all other distinct platelet variants on chromosome 6. Additionally, these results link the variants to putative target genes via our gene expression prediction models.

Figure 4 shows that there is cell-type specific epigenetic evidence that supports our findings. *IRAK1BP1* is a component of the IRAK1-dependent TNFRSF1A signaling pathway, which can activate NF-kappa-B and regulate cellular apoptosis and inflammation.^27^ Variants in the gene expression prediction model for *IRAK1BP1* in high LD with chr6:79617522 overlapped with megakaryocyte ATACseq peaks from BLUEPRINT (Figure 4e).^24^ Additionally, we observed via megakaryocyte pcHi-C data that these same variants in the *IRAK1BP1* prediction model interact with the promoter region for the nearby gene, lebercilin LCA5 (*LCA5*) (Figure 4c). *LCA5* plays roles in centrosomal functions in nonciliary cells.^28^ While both *IRAK1BP1* and *LCA5* are expressed in megakaryocyte cells using expression data from BLUEPRINT, the expression level is higher in *LCA5*, suggesting a potential role for *LCA5* in platelet trait variability, despite not being captured by TWAS (Figure 4d). *LCA5* is not present in the DGN reference panel, and thus unavailable to fit a prediction model, likely because of low expression in whole blood (median TPM 0.018 in GTEx v8).^29^ Integration of our TWAS results with expression and chromatin conformation data in platelet producing megakaryocyte cells reveals novel candidate genes at this genomic locus; it is possible that the variants in *IRAK1BP1* aggregated by the TWAS prediction model impact the expression of *LCA5* through spatial proximity to the promoter region of the gene. The *IRAK1BP1* locus shows both the statistical power gains possible through TWAS analysis and the importance of full consideration of other potential target genes as well as complementary functional annotation resources in biological interpretation of a TWAS identified signal.

#### TWAS Genes Implicated in Novel Phenotype Categories

Our TWAS conditional analysis identified 92 conditionally significant associations grouped into 70 loci with no distinct GWAS sentinels for the corresponding phenotype category within 1Mb of the gene. These gene-trait associations represent novel TWAS findings; our results support that the previously reported association at the locus is extended to a new class of correlated phenotypes (for example, extension of loci already associated with red blood cell related traits to platelet or white blood cell indices). Among the 92 associations, 42 were unable to be assessed for replication in MVP due to unavailable phenotype data or missing genotypes for at least one variant in the TWAS prediction model, and 35 out of the remaining 50 were replicated at a nominal significance threshold (*α* = 0.05) with the same direction of effect as in UKB. 17 out of 50 are replicated at the Bonferroni adjusted threshold for the total number of conditionally significant associations (*α* = 0.05/557 = 8.98×10^-5^).

##### *CD79B* (GRCh37 chr17:62,006,100 - 62,009,714)

One such example is the 1Mb region surrounding B-cell antigen receptor complex-associated protein beta chain (*CD79B*), which was associated with lymphocyte count (p = 9.81×10^-10^), hematocrit (p = 1.22×10^-9^), plateletcrit (p = 3.37×10^-9^), white blood cell count (p = 8.49×10^-9^), and hemoglobin percentage (p = 1.21×10^-7^) in our TWAS marginal analysis. Supporting the role of this gene in blood cell indices, an extremely rare mutation in *CD79B*, rs267606711, has been reported to cause agammaglobulinemia 6 [MIM: 612692], an immunodeficiency characterized by profoundly low or absent serum antibodies and low or absent circulating B cells due to an early block of B-cell development^30, 31^.

In *Vuckovic et al.*, the region surrounding *CD79B* contained several borderline genome-wide significant variants for lymphocyte count, with lead variant 17:62008437_C_T (p = 2.3×10^-9^). However, in their conditional analysis, the region was clumped into nearby lymphocyte count GWAS signals, namely 17:57929535_A_G (p = 1.16×10^-25^) with annotated target gene RNA, U6 small nuclear 450, pseudogene (*RNU6-450P*) and 17:65087308_G_C (p = 4.34×10^-10^) with target gene helicase with zinc finger (*HELZ*) (with both genes assigned based on distance). After conditioning on the set of 186 white blood cell count distinct variants identified by GWAS conditional analysis on chromosome 17, including 17:57929535_A_G and 17:65087308_G_C, *CD79B* continued to demonstrate evidence of association with lymphocyte count (p = 9.8×10^-10^) and white blood cell count (p = 8.5×10^-9^).

Further, there were 6 distinct GWAS signals from individual level GWAS conditional analysis across both red blood cell and platelet traits within 1Mb of *CD79B*. To control for confounding due to correlated hematological traits, we further conditioned on the 6 distinct variants for red blood cell and platelet traits in addition to the set of 186 white blood cell distinct variants. The association with lymphocyte count remained nominally significant (p = 3.03×10^-4^) and the white blood cell count association was attenuated (p = 0.16). *CD79B* demonstrated some evidence of association with lymphocyte count in MVP Europeans as well (p = 1.1×10^-5^) with matching direction of association, despite not achieving the Bonferroni adjusted threshold (*α* = 8.34×10^-6^). Therefore, supporting the increased power of TWAS above single variant GWAS, our findings suggest the biologically plausible *CD79B* association with lymphocyte count was likely distinct from previously reported genetic loci in the neighborhood.

### TWAS fine mapping via Conditional Analysis

TWAS conditional analysis was also used to fine map TWAS loci in which multiple genes achieved the Bonferroni adjusted significance threshold (see Supplemental Table S2). For example, erythropoietin (*EPO*) (GRCh37 chr7:100,720,800-100,723,700) encodes the primary regulator of red blood cell production and has been well studied for its impact on blood cells through its causal role in familial erythrocytosis [MIM: 617907] and Diamond-Blackfan anemia-like [MIM: 617911].^32^ The 1Mb region surrounding *EPO* also contains 28 distinct GWAS signals across 21 different blood cell traits. In our marginal TWAS analysis, 13 marginally significant TWAS genes associated with hemoglobin concentration at this locus including *EPO* (p = 2×10^-12^), with the TWAS sentinel gene being solute carrier family 12 member 9 (*SLC12A9*) (p = 2.51×10^-29^). However, despite the well-studied links to blood cell genetics, the *EPO* gene was not included in the 95% FINEMAP credible set. Yet after we condition the TWAS predicted expression on the distinct red blood cell signals at this locus, *EPO* was the only conditionally significant gene at the locus (p = 2.19×10^-6^). The association between *SLC12A9* and hemoglobin was completely attenuated after conditioning (p = 0.71). This suggests that while the genetic link between *EPO* and blood cell traits are well established, the full set of causal variants and overall genetic architecture underlying the association remains elusive.

### TWAS-based assignment of variants to target genes

To compare co-localization and TWAS approaches of assigning GWAS variants to potential causal genes, we considered 10,239 variant-trait associations across 10 hematological traits from Vuckovic *et al*. (see **Methods**). In their analyses, *coloc* identified 427 out of 10,239 associations (4.2%) which colocalized with at least one eQTL. (Figure 5a). We take the eGene(s) corresponding to these eQTLs as the gene(s) assigned to the GWAS variant by *coloc.* In comparison, our TWAS based approach assigned target genes to 1,738 variant-trait associations, a four-fold increase compared to *coloc*. Of the 269 associations assigned to at least one gene by both methods, 80% of the associations have at least one assigned gene in common, demonstrating that the two methods tend to assign variants to the same genes where they both assign a target gene. Of the 158 associations assigned to genes by *coloc* but not by our TWAS-based approach, 13 were assigned to genes with no expression data in our DGN reference dataset, 23 were assigned to TWAS genes with poor model predictive performance (model R2 <= 0.05), 51 variant-trait associations were not within +/-1MB of any TWAS loci, 49 were only nearby TWAS loci with a non-significant sentinel gene, and 22 had low correlations between variant dosage for the lead GWAS variant and imputed TWAS gene expression (r2 < 0.2) (Figure S6).

**Figure 5.**
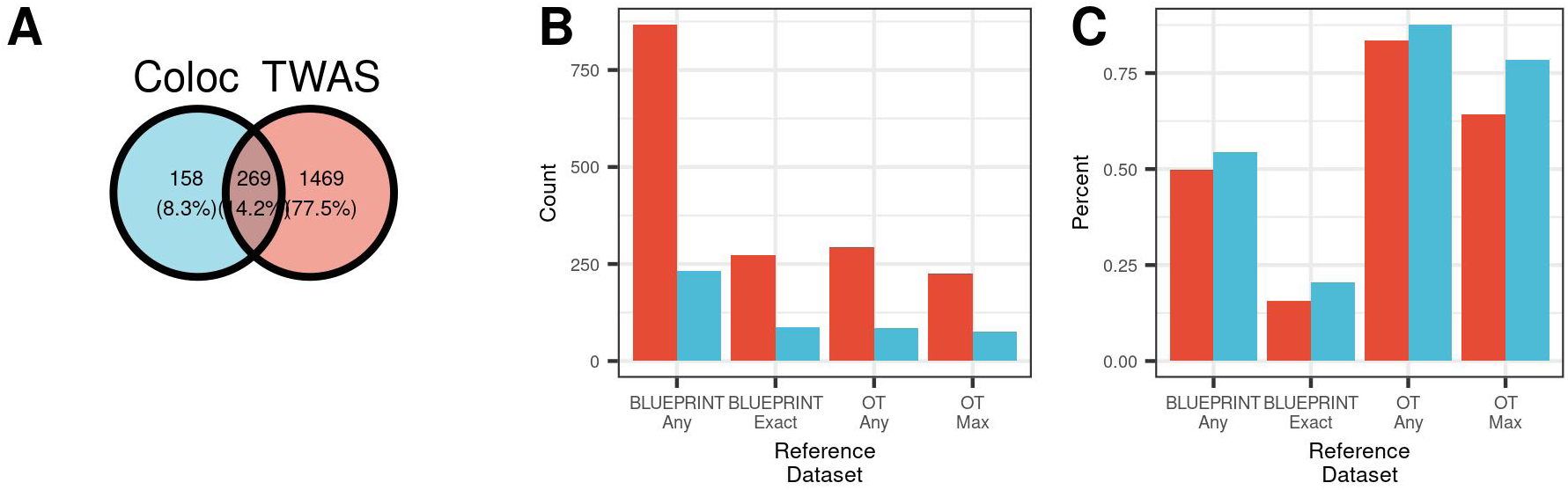
TWAS and coloc variant-to-gene assignments. We compare our TWAS-based variant-to-gene assignments with assignments from *coloc* using a set of 10,239 variants associated with 10 hematological traits. (A) *Coloc* successfully assigns 427 variants to target causal genes, while our TWAS based approach assigns 1,738 to target genes. (B-C) We compare these assignments in a variety of settings, using variant-to-gene assignments both considering phenotype-specific and phenotype-agnostic approaches. The TWAS based approach has increased sensitivity to assign genes to potentially causal genes (B) and decreased specificity to *coloc* (C).

To illustrate one example where the two methods agree, Figure 6 highlights the concordant TWAS and *coloc* assignment of rs6062304 (GRCh37 chr20:62351539_A_T), a distinct variant for lymphocyte percentage, to Lck interacting transmembrane adaptor 1 (*LIME1*), a gene with known involvement in T cell signaling.^33,34^ In Vuckovic *et al.*, rs6062304 was assigned via VEP annotation to zinc finger CCCH-type and G-patch domain containing (*ZGPAT*), which has no clear link to blood cells. Figure 6a shows Vuckovic *et al.* GWAS results overlaid with the marginal TWAS results for lymphocyte percentage. Six TWAS genes are significant, and a subset of 3 genes are included in the FINEMAP 95% credible set: *LIME1, ZGPAT,* and regulator of telomere elongation helicase 1 (*RTEL1*). Figure 6a shows that *LIME1* predicted expression is highly correlated (r2 = 0.905) with rs6062304, while *ZGPAT* is moderately correlated (r2 = 0.556). *coloc* assigns *LIME1* as an eGene because of the high LD (r2 = 0.916) between rs6062304 and an eQTL for *LIME1*, rs6062497 (Figure 6b). Similarly, Figure 6c demonstrates that variants with the largest weights in the *LIME1* gene expression prediction model are in high LD with rs6062304. In contrast, Figure 6d reveals that variants in high LD with rs6062304 have smaller TWAS weights in the *ZGPAT* model, suggesting that the *ZGPAT* association with lymphocytes at this locus is not primarily due to rs6062304. While both *LIME1* and *ZGPAT* correlations pass the r2 cutoff for the TWAS-based gene assignment (r2 > 0.2), *LIME1* predicted expression is much more correlated with rs6062304, and is the most likely target gene at this locus according to the TWAS based approach. This highlights the value of considering correlation of predicted gene expression with the lead GWAS variant in TWAS assignment of likely target genes, as done in our pipeline. Thus, using different approaches, TWAS-based and *coloc*-based variant-to-gene assignment methods assign rs6062304 to a biologically plausible target gene, improving upon distance-based approaches.

**Figure 6.**
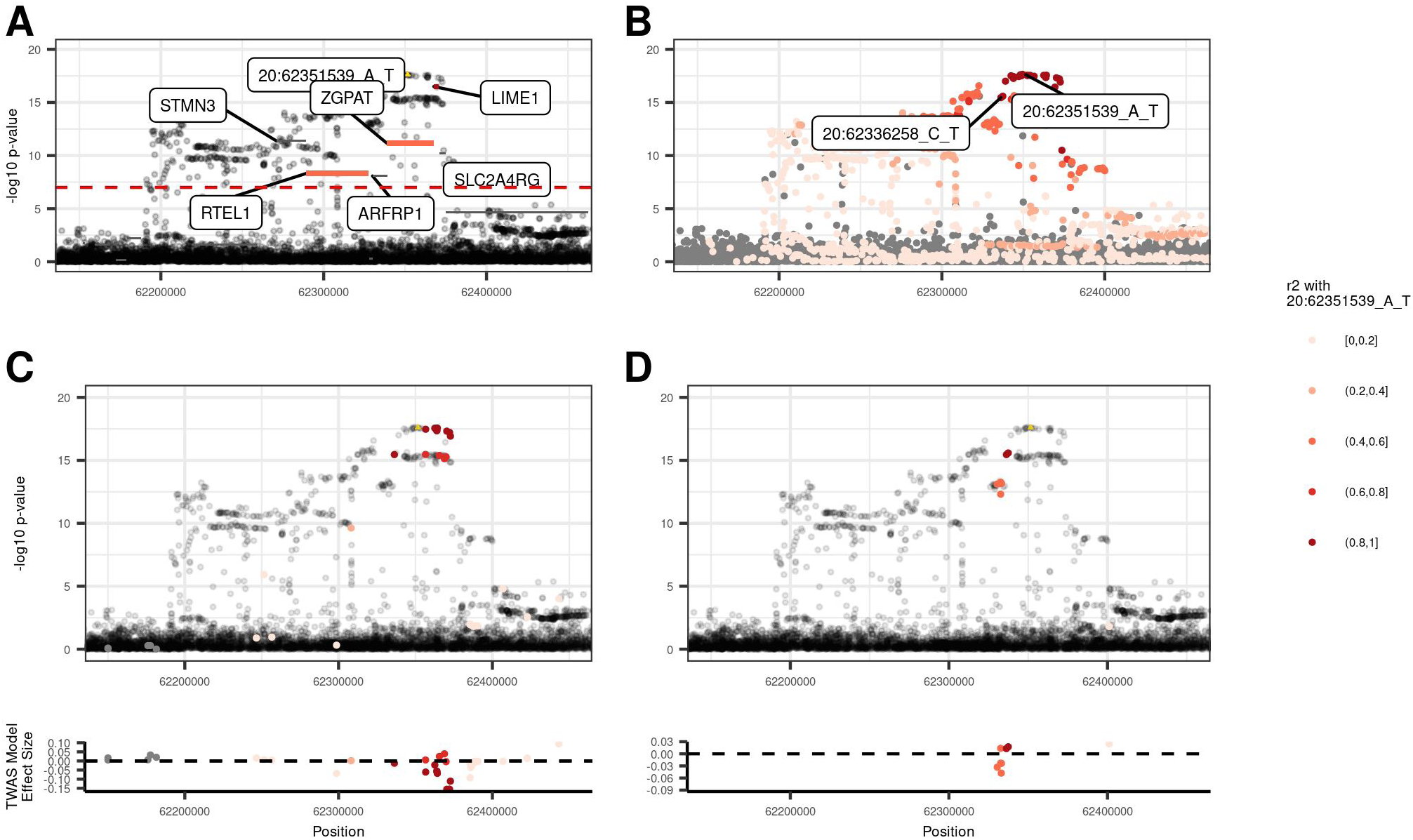
TWAS and *coloc* variant-to-gene assignments agree at *LIME1* locus. The *LIME1-*lymphocyte percentage associated locus illustrates one example where the TWAS- and *coloc-* based variant-to-gene assignments agree. (A) Genes in the FINEMAP credible set are colored by their correlation with rs6062304. The predicted gene expression with *LIME1* is highly correlated with rs6062304, whereas neither *ZGPAT* nor *RTEL1* are. (B) An eQTL for *LIME1*, rs6062497, is in high LD with rs6062304, and in turn *coloc* assigns *LIME1* as an eGene. (C-D) Model variants for *LIME1* and *ZGPAT* are colored by their LD with rs6062304, respectively. The variants with the largest effect sizes in the TWAS prediction model for *LIME1* are in high LD with rs6062304, whereas those for *ZGPAT* are not.

However, in scenarios where eQTLs have not been identified in a target tissue of interest, likely due to small sample size for a given expression dataset, TWAS-based methods, which combine multiple potential eQTLs which may be in LD with a GWAS variant, are better powered to assign GWAS variants to target genes. Figure 5a shows that there are 1,469 variant-trait pairs which are assigned to a target gene via TWAS not assigned to a gene by *coloc*. One such example is the TWAS assignment of rs1985157 to leucine rich repeat containing 25 (*LRRC25*) (GRCh37 chr19:18513594_T_C), a distinct variant for neutrophil count and neutrophil percentage. Neither VEP nor *coloc* assigned rs1985157 to a target gene. Our TWAS marginal analysis identified 4 significant genes for neutrophil count at this locus, *LRRC25,* elongation factor for RNA polymerase II (*ELL*), single stranded DNA binding protein 4 (*SSBP4*), and inositol-3-phosphate synthase 1 (*ISYNA1*) (Figure 7a). However, only *LRRC25* predicted gene expression values have a strong correlation with rs1985157 (r2 = 0.863). Two other TWAS-assigned genes are moderately correlated with rs1985157 (*ELL* r2 = 0.46) and (*SSBP4* r2 = 0.47). Figure 7c shows that variants in the *LRRC25* prediction model that are in high LD with rs1985157 have the largest weights in absolute value. In contrast, Figure 7d shows that *SSBP4* predicted expression is driven by variants in moderate LD with rs1985157. Several studies have suggested that *LRRC25* plays a key role in innate immune response and autophagy.^35, 36^ Further, cell-type specific gene expression data from BLUEPRINT suggest that *LRRC25* is specifically expressed in neutrophils.^24^ Our results show that TWAS-based variant-to-gene assignment methods can identify biologically plausible target genes, even when *coloc* fails to do so.

**Figure 7.**
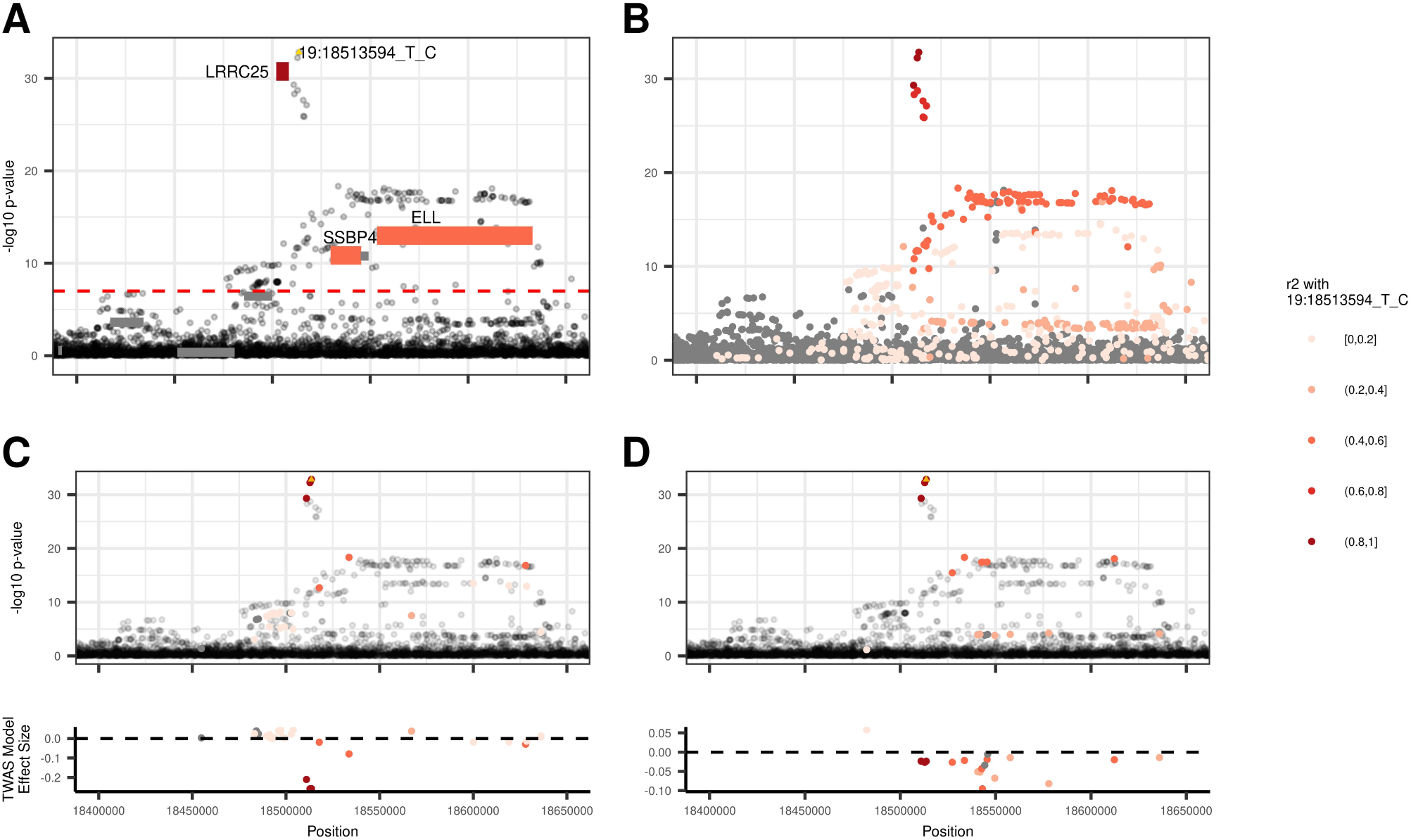
TWAS assigns rs1985157 to *LRRC25*. Figure 7 illustrates how TWAS assigned rs1985157 to *LRRC25* when *coloc* failed to do so with no eQTL in the region. (A) *LRRC25* predicted gene expression was highly correlated with rs1985157 (r2 = 0.863), whereas the prediction from *SSBP4* (r2 = 0.47) and *ELL* (r2 = 0.457) were not as highly correlated despite both genes being significant. (B) LD patterns for variants at the locus. (C-D) Only model variants for *LRRC25* and *SSBP4* are colored by their LD with rs1985157, respectively. Variants with the largest TWAS weights (in absolute values) for *LRRC25* are in high LD with rs1985157, whereas those for *SSBP4* are not.

In order to understand the differences in the TWAS and *coloc* gene assignments and to examine whether the additional variants assigned to genes by TWAS over *coloc* have relevant epigenetic evidence to the phenotype of interest, we compared the gene assignments of TWAS and *coloc* using BLUEPRINT cell-type specific expression data and Open Targets V2G scores (see **Methods** for details).^23^ We hypothesized that genes with cell-type specific expression in BLUEPRINT in relevant cell types are more likely to be relevant target genes for GWAS associations.

We found that the TWAS-based approach assigned GWAS variants to genes identified by external datasets at a slightly lower rate than the *coloc* assignments, but identified target genes for more than double the number of variants (see Figure 5b-c). Specifically, Figure 5d shows that 84% of the TWAS-based variant-to-gene assignments are supported by Open Targets (OT Any genes), and 64% of genes assigned by TWAS are the most likely target gene as identified by Open Targets (OT Max gene). In comparison, 88% of the *coloc* assigned genes are supported by OT Any genes and 78% as the OT Max gene. On the other hand, Figure 5c shows that 294 TWAS pairs are assigned to an OT Any gene and 226 pairs assigned to an OT Max gene, much larger number of supported assignments than the 85 and 76 *coloc* pairs, a 3.46 and 1.97 fold increase respectively. The proportion of variants assigned to cell-type specifically expressed genes in BLUEPRINT expression data is lower compared to the Open Targets assignments (Figure 5d). However, the TWAS-based approach matches 3.12-fold more variants to specifically expressed genes in trait-relevant cell types and 3.73-fold more genes to specifically expressed genes for any blood cell compared to *coloc*. Therefore, our results suggest that TWAS, compared to *coloc*, is less specific but more sensitive when assigning variants to target genes.

We then applied our TWAS-based variant-to-gene assignment to all 29 hematological traits considered in our UKB TWAS. We successfully assigned 4,261 variant-trait associations to 1,842 distinct potentially causal TWAS genes with an average of 1.45 (SD = 0.81) genes assigned per variant-trait association (see Supplemental Table S3). Of the 4,261 associations, 746 (17.5%) were assigned to specifically expressed genes in trait-relevant cell types, and 1,982 (46.5%) were assigned to specifically expressed genes for any blood cell. Both rates were comparable to the performance of the TWAS variant-to-gene assignments in the phenotype restricted results above. For the 813 overlapping variant-to-gene assignments from the Open Targets datasets, the replication rates were similar to the phenotype restricted results for OT Any genes (78.2%), but the replication rate decreased for the OT Max gene (54.5%).

## Discussion

Our TWAS of blood cell traits in UKB Europeans demonstrates the utility of TWAS to identify novel loci and to extend known loci to additional phenotype categories, even in well-studied hematological traits for which over 10,000 loci have been reported by previous GWAS studies^2,3^.

In total, our TWAS conditional analysis identifies 557 conditionally significant TWAS loci across 29 blood cell phenotypes, 199 of which replicated in MVP at a nominal significance level and 108 at a Bonferroni adjusted threshold. Our TWAS conditional analysis results suggest that as GWAS sample sizes and statistical power for single variant analysis continue to increase, additional statistically distinct variant signals, with lower allele frequencies or effect sizes, will continue to be discovered. Often, these conditionally significant TWAS genes are not the most marginally significant genes at their respective TWAS locus, suggesting that marginal TWAS results can be driven by previously discovered GWAS variants. Despite the promises of TWAS offering increased interpretability for GWAS results, deciphering the relationship between variant and gene level effects remains challenging.^20, 37, 38^

As a subset of the 557 conditionally significant results, we identify 10 novel loci previously undiscovered even after several large-scale GWAS in both European and trans-ethnic cohorts. In addition, we replicate 3 out of 6 of the novel TWAS loci available for replication in MVP at a nominal significance threshold, and 1 at a Bonferroni adjusted significance threshold. Used in a less studied complex trait compared to hematological traits, a marginal TWAS analysis may be expected to identify even more novel signals, due to enhanced power over single variant GWAS analyses. For traits with a large sample size GWAS previously conducted, combining TWAS conditional analysis with previously conducted GWAS conditional analysis can further expand our ability to distinguish statistically distinct signals and link variants to target genes at complex loci, as demonstrated by our analysis at the *IRAK1BP1* locus.

Further, we demonstrate the utility of TWAS conditional analysis to extend previously reported associations at genomic loci to a new class of correlated phenotypes, identifying 92 such loci. Due to the shared genetic architecture of blood cell traits which is mediated through the differentiating of common progenitor cells, variants which impact one class of blood cell traits may plausibly have an effect on other hematological traits. However, through conditioning our TWAS predicted expression on all distinct blood cell associated variants in a region, we demonstrate that TWAS can identify loci associations which are statistically distinct from previous GWAS discoveries. The *CD79B* locus demonstrates a robust association with lymphocyte count despite conditioning on previously identified white blood cell, red blood cell and platelet associated variants at the locus. This robust association confirms previously reported biological roles for *CD79B* with lymphocyte function, and establishes relevant variant-level candidates for functional validation through the TWAS prediction model.^30, 31^

We address some challenges with interpreting TWAS loci at the scale of biobank-sized analyses through our adapted TWAS fine mapping via FINEMAP and conditional analysis with individual level TWAS data.^20^ To our knowledge, all TWAS fine mapping methods and software are currently designed for summary-statistics based TWAS approaches, with limited functionality to input user generated TWAS statistics from individual level data such as those generated by our REGENIE-based approach.^37, 39^ To overcome this challenge, we substitute the variant-level LD matrix for the predicted expression correlation matrix in our UKB sample in the widely-used FINEMAP software to generate credible sets of genes. However, the main shortcoming of our approach is that we are only addressing correlation at the gene level via predicted expression values. In addition to previous research, future methodological research and software development should be done to address this challenge.^37, 38^

Our application of TWAS conditional analysis to TWAS locus fine mapping as demonstrated through the example of the *EPO* locus provides a way forward to combine trait-specific variant-level and gene-level information to identify gene-trait associations which are not driven by existing GWAS knowledge, where additional single variant signals likely remain to be discovered. This approach could be extended to consider step-wise TWAS-on-TWAS conditional analyses to generate credible sets of genes at a locus, similar to step-wise analyses in GWAS based on individual level data. Direct conditional analysis is only possible when using individual level genotype data in the discovery cohort, which presents an advantage of using non-summary statistics based methods to perform TWAS when possible. In addition, we are better able to account for confounding due to LD from a potentially mis-specified LD reference panel than when using summary statistics based methods for both TWAS and conditional analysis. While improving the match of LD reference panels to study populations, especially for non-European populations, will improve the usefulness of summary statistics based TWAS, our work suggests distinct advantages of leveraging individual level genotype data when possible.

Additionally, we perform systematic variant-to-gene assignment for distinct hematological trait GWAS signals using a TWAS-based approach, and demonstrate that many of our assignments are supported by external resources. Our results suggest that our TWAS-based approach of assigning GWAS variants to target genes can map many more variants to target genes using biobank scale data. This increased number of variants assigned to target genes comes at a price of decreased sensitivity. At both the *LRR*C25 and *LIME1* loci, TWAS identifies additional genes that are moderately correlated with the GWAS variant of interest. Thus, our results support complementary roles of TWAS and colocalization approaches. We propose first using colocalization to assign GWAS variants to target genes using available cell-type specific eQTL data relevant to the trait of interest, and then leveraging the additional assignments generated by TWAS for GWAS variants not assigned to a target gene. Additionally, several refinements could be made to the TWAS-based variant-to-gene assignments. By leveraging cell/tissue type specific gene expression datasets to train gene expression models in the future, the TWAS based approach could be extended to match the common practice of conducting eQTL colocalization or other target gene assignment analyses in trait-relevant cell/tissue types.^40–43^ The TWAS based approach could be further refined through a modified conditional TWAS analysis, in which a GWAS variant is conditioned separately on predicted gene expression values of potential target genes at the locus, and the attenuation in the GWAS signal for each gene is assessed.

There are still several future directions for the improvement of biobank-scale TWAS studies. First, increasing sample sizes in tissue specific expression datasets will allow future TWAS studies to train gene expression prediction models in cell/tissue types which are directly relevant to tissues of interest. Already, several TWAS methods have been developed to leverage multiple tissues to train better gene expression prediction models.^6,7, 44^ Additionally, the TWAS variant-to-gene assignment approach would benefit from larger expression datasets to train cell/tissue type specific gene expression prediction models to assess the correlation between predicted expression and a GWAS variant of interest across several relevant models. For example, at the identified *IRAK1BP1* locus, it would be useful to have larger megakaryocyte specific gene expression datasets available for TWAS model training; similar cell-type specific panels would be useful for other hematological indices and for complex trait analysis more generally. Such cell-type specific reference panels are becoming increasingly available, though not always in adequate sample sizes for TWAS and not always with publicly available individual level data.^45^ Second, extending the variable selection procedure for prediction models past the 1Mb *cis*-region surrounding TWAS genes either via trans-eQTL datasets or by selecting variants which are highly likely to be in interesting epigenomic regions will improve TWAS models.^46, 47^ In summary, we conduct a large-scale TWAS of well-studied hematological traits and discover novel loci. We show that TWAS-based approaches for assigning variants to their target genes are comparable in specificity to co-localization based approaches, but are able to assign many more variants (4.07 fold increase) to target genes. Our careful use of conditional analysis, TWAS-based fine mapping, and TWAS-based variant-to-gene assignments in the context of blood cell traits will be broadly useful to the practice of TWAS for other complex traits.

## Supplemental Information description

Supplemental information contains 3 tables and 6 figures.

## Acknowledgements

B.R. was supported by the National Science Foundation Graduate Research Fellowship Program under Grant No. DGN-1650116. V.S. was supported by National Institutes of Health (NIH) R01DK103794. G.V. was supported by K08MH122911, NIMH; BBRF Young Investigator Grant #29350, BBRF. This study was also supported by the National Institutes of Health (NIH), Bethesda, MD under award numbers R01AG050986, R01AG065582, R01AG067025, U01MH116442, R01MH109677 (P.R.), and by the Veterans Affairs Merit grants BX002395 and BX004189 (P.R.). The project described was supported by the National Center for Advancing Translational Sciences, National Institutes of Health, through Grant KL2TR002490 (LMR). Y.L. was supported by NIH R01 HL129132; R01 HL146500; R01 GM105785.

This research has been conducted using the UK Biobank Resource under Application Number 25953. Support for title page creation and format was provided by AuthorArranger, a tool developed at the National Cancer Institute.

This research is based on data from the Million Veteran Program, Office of Research and Development, Veterans Health Administration, and was supported by awards #MVP006 and #MVP001. This publication does not represent the views of the Department of Veteran Affairs or the United States Government.

The Genotype-Tissue Expression (GTEx) Project was supported by the Common Fund of the Office of the Director of the National Institutes of Health, and by NCI, NHGRI, NHLBI, NIDA, NIMH, and NINDS. The data used for the analyses described in this manuscript were obtained from: the GTEx Portal on 05/01/2021. Trans-Omics in Precision Medicine (TOPMed) program imputation panel (version TOPMed-r2) supported by the National Heart, Lung and Blood Institute (NHLBI); see www.nhlbiwgs.org. TOPMed study investigators contributed data to the reference panel, which can be accessed through the Michigan Imputation Server; see https://imputationserver.sph.umich.edu. The panel was constructed and implemented by the TOPMed Informatics Research Center at the University of Michigan (3R01HL-117626-02S1; contract HHSN268201800002I). The TOPMed Data Coordinating Center (3R01HL-120393-02S1; contract HHSN268201800001I) provided additional data management, sample identity checks, and overall program coordination and support. We gratefully acknowledge the studies and participants who provided biological samples and data for TOPMed.

## Web Resources

TOPMed Imputation Server: https://imputation.biodatacatalyst.nhlbi.nih.gov/#

REGENIE: https://rgcgithub.github.io/regenie/

boltLMM: https://alkesgroup.broadinstitute.org/BOLT-LMM/BOLT-LMM_manual.html

FINEMAP: http://www.christianbenner.com/#

GTEx Portal: https://www.gtexportal.org/home/

OMIM: https://www.omim.org/

WashU Epigenome Browser: https://epigenomegateway.wustl.edu/

TOP-LD: http://topld.genetics.unc.edu/topld/

## Data and code availability

This paper did not generate any new datasets. Codes to analyze the data can be found at https://github.com/brycerowland/UKB_BCT_TWAS.git.

