## Supplemental Figures and Methods for "Transcriptome-wide association study in UK Biobank Europeans identifies associations with blood cell traits"

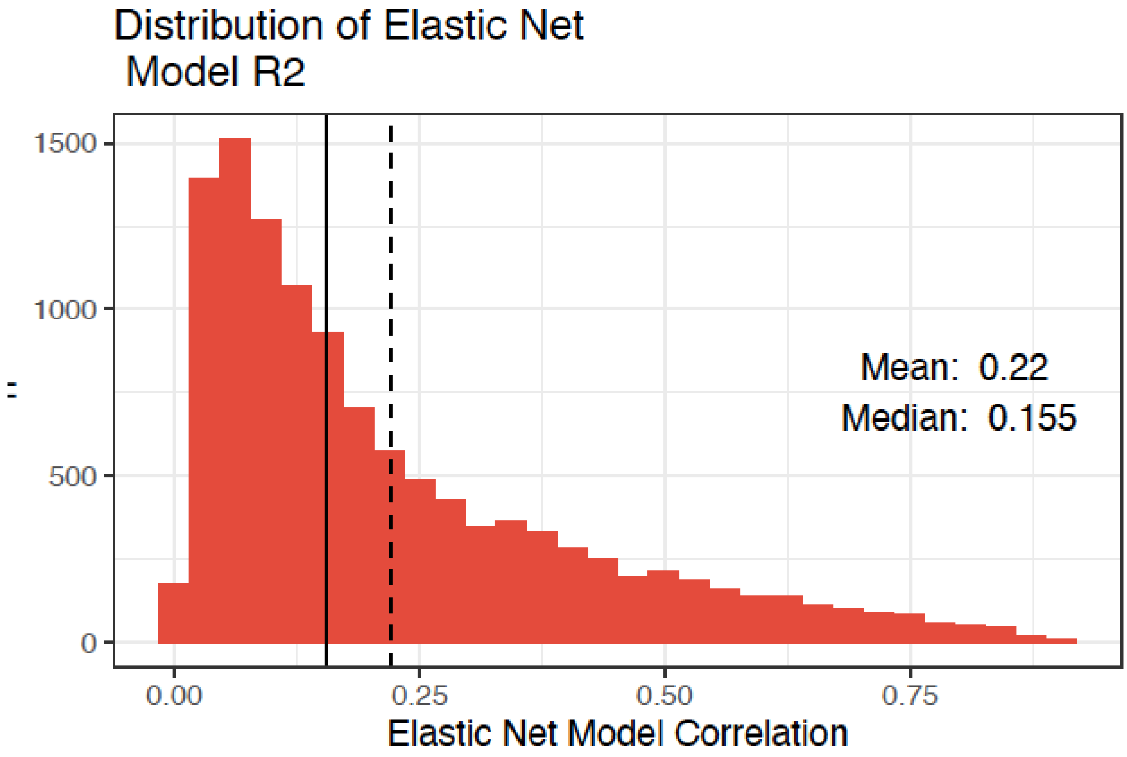


***Figure S1*** **-** Distribution of Model R2 of gene expression prediction models trained with Elastic Net.

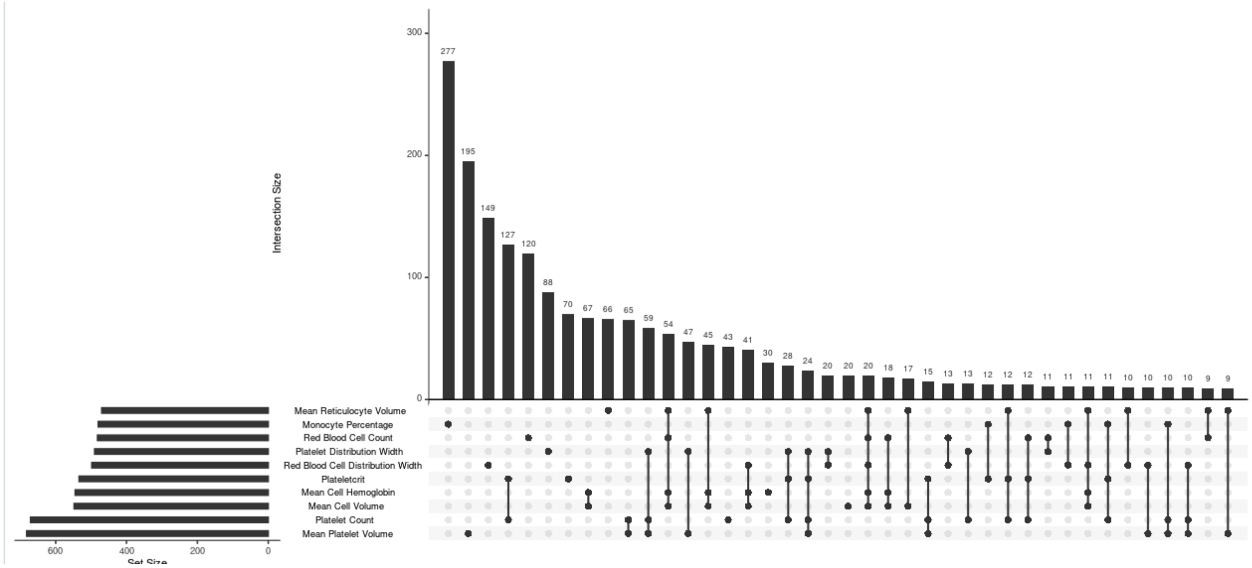


***Figure S2*** **-** Upset Plot of marginal analysis genes by phenotype.

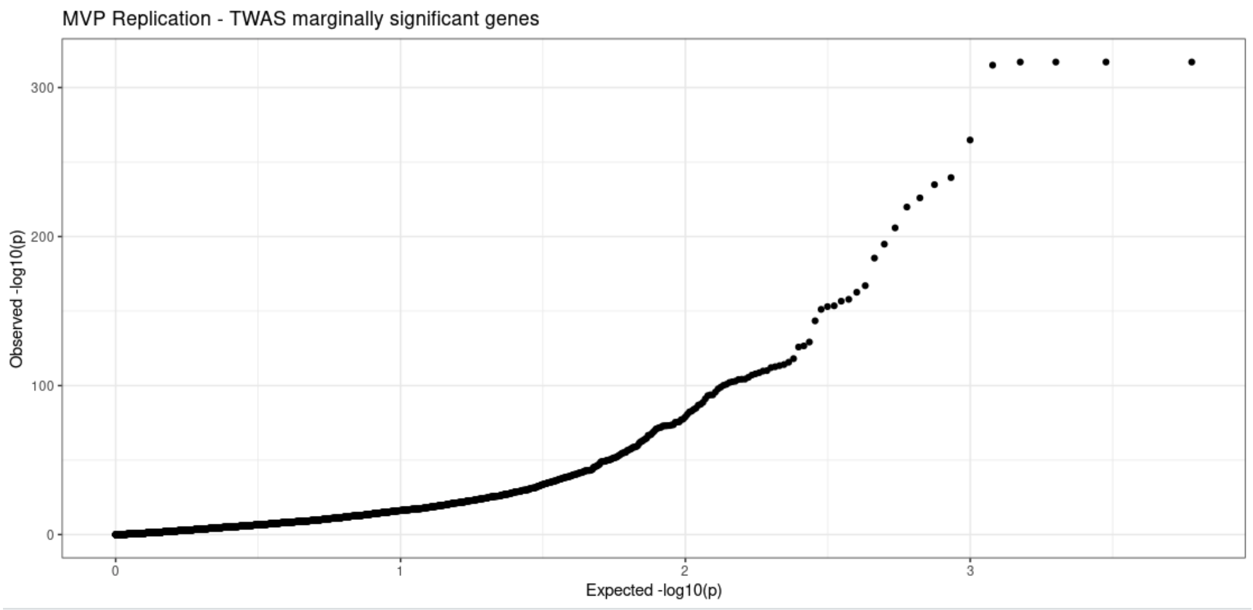


***Figure S3*** - QQPlot of -log10 p-values from MVP Replication of marginally significant TWAS genes.

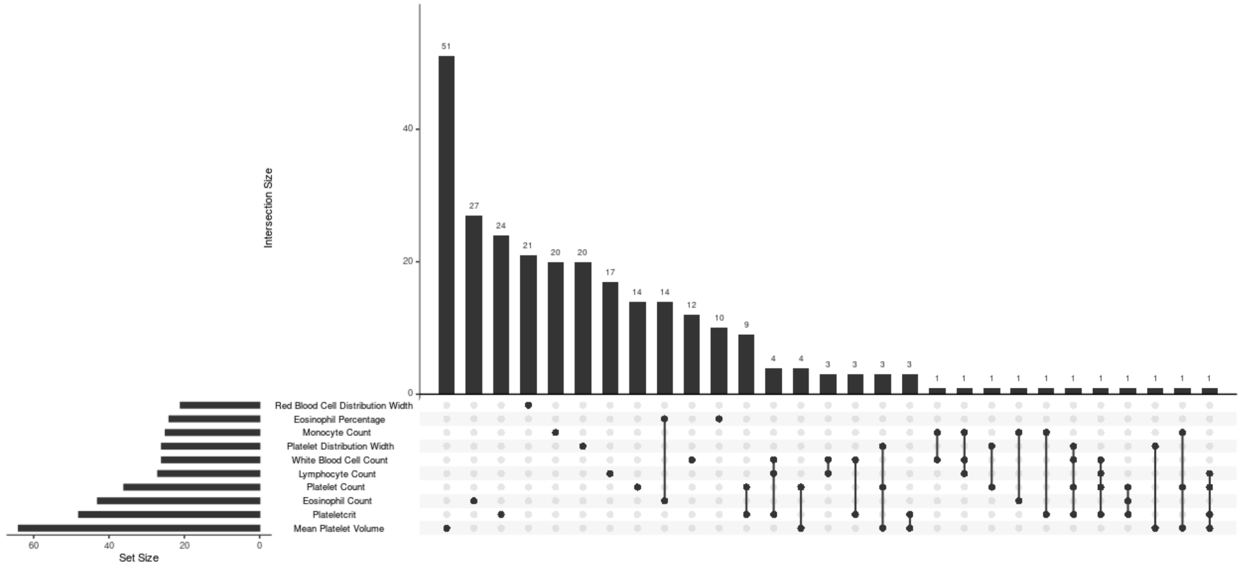


***Figure S4*** - Upset Plot of conditional analysis genes by phenotype.

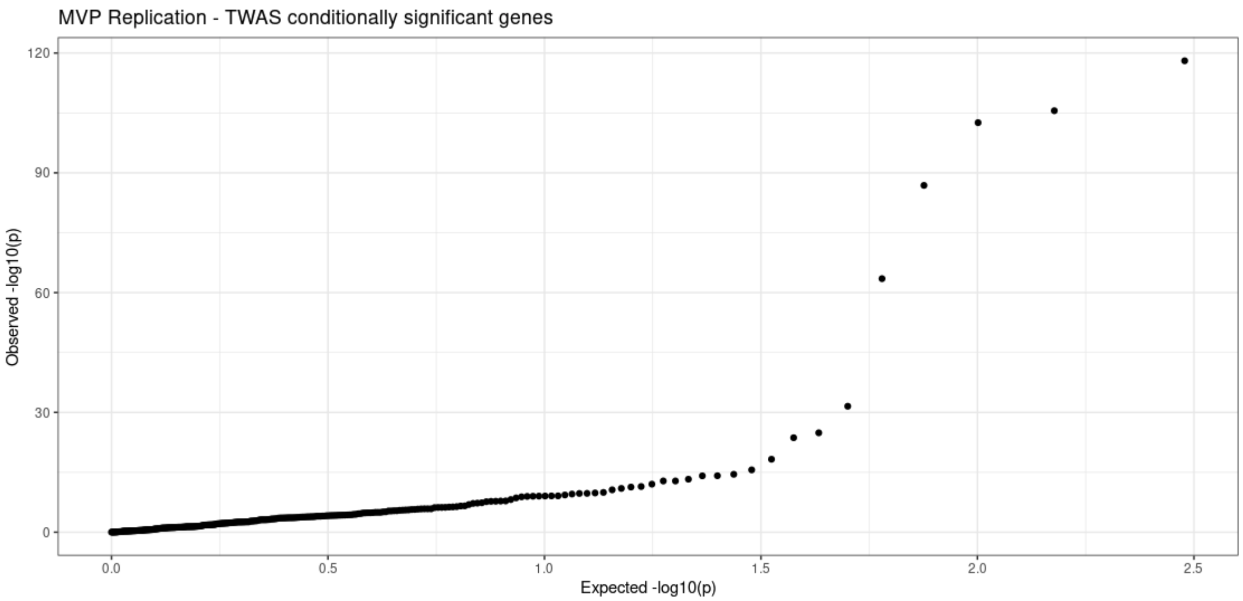


***Figure S5*** **-** QQPlot of -log10 p-values from MVP Replication of conditionally significant TWAS genes.

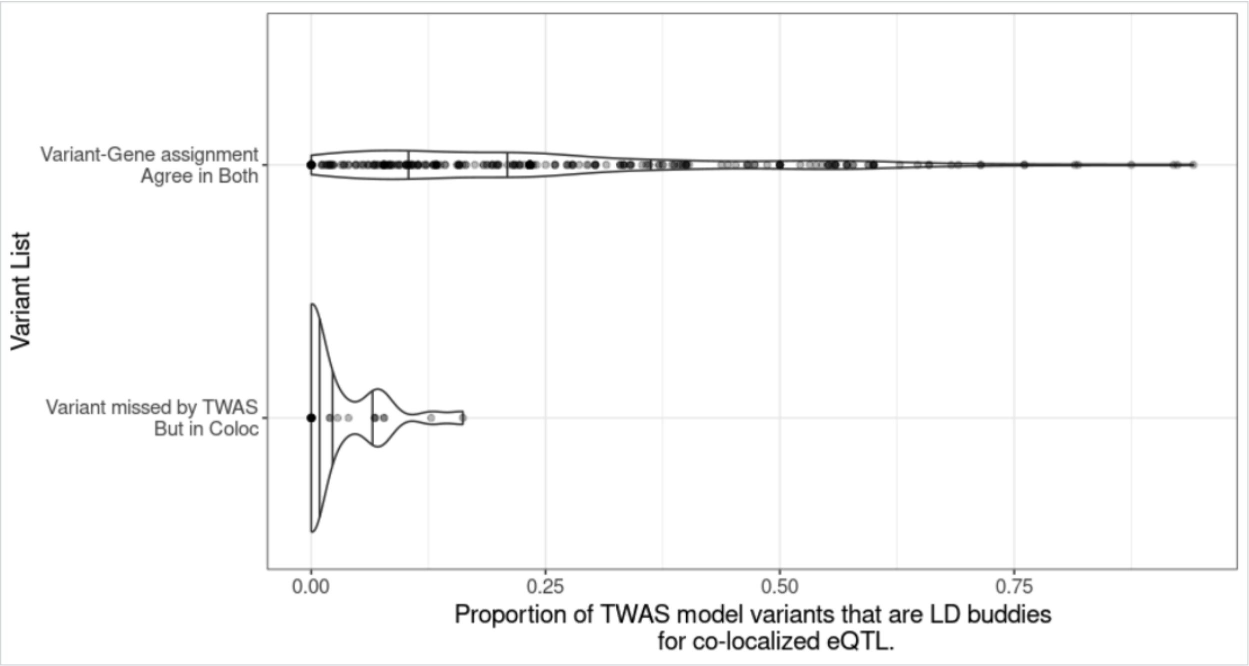


***Figure S6*** **- Variant assignments missed by TWAS but identified by *coloc* are driven by variants independent of co-localized eQTL.** We compared the 22 associations where the TWAS-based approach fails to assign target genes despite *coloc* identifying these variants as co-localized with an eQTL for a gene in the region to the variant-trait associations assigned by both methods. Supplemental Figure 6 shows that variants assigned to target genes by *coloc* but not by TWAS were more likely to co-localize to eQTLs which are not represented in the TWAS gene expression prediction model. The proportion of variants in LD (1000G EUR r2>0.5) with the eQTL which co-localized with the GWAS sentinel variant in the TWAS prediction model was remarkably lower than the proportions for variant-gene pairs identified by both methods. This suggests that in these 22 prediction models, which were TWAS-significantly associated with the phenotype of interest, the TWAS predictions are not driven by the co-localized eQTLs, but other variants at the locus.

### Supplemental Methods

#### ICD9/10 Exclusions

**ICD9**: 170, 1700, 1701, 1702, 1703, 1704, 1705, 1706, 1707, 1708, 1709, 200, 2000, 2001, 2002, 2008, 201, 2010, 2011, 2012, 2014, 2015, 2016, 2017, 2019, 202, 2020, 2021, 2022, 2023, 2024, 2025, 2026, 2028, 2029, 203, 2030, 2031, 2038, 204, 2040, 2041, 2042, 2048, 2049, 205, 2050, 2051, 2052, 2053, 2058, 2059, 206, 2060, 2061, 2062, 2068, 2069, 207, 2070, 2071, 2072, 2078, 208, 2080, 2081, 2082, 2088, 2089, 2384, 2385, 2386, 2387, 282, 2820, 2821, 2822, 28220, 28221, 28222, 28229, 2823, 2824, 28240, 28241, 28242, 28243, 28244, 28245, 28246, 28247, 28248, 28249, 2825, 2826, 2827, 28270, 28271, 28272, 28273, 28274, 28275, 28279, 2828, 2829, 283, 2830, 2831, 28310, 28311, 28312, 28313, 28314, 28315, 28319, 2832, 28320, 28321, 28322, 28329, 2839, 28390, 28391, 28399, 284, 2840, 28400, 28401, 28402, 28409, 2848, 28480, 28481, 28482, 28483, 28484, 28485, 28489, 2849, 286, 2860, 2861, 2862, 2863, 28630, 28639, 2864, 2865, 2866, 2867, 28670, 28679, 2869, 287, 2870, 2871, 2872, 2873, 28730, 28731, 28732, 28739, 2874, 28740, 28741, 28742, 28749, 2875, 2878, 2879, 288, 2880, 28800, 28801, 28802, 28803, 28804, 28809, 2881, 2882, 2883, 2888, 28880, 28881, 28889, 2889, 289, 2890, 28900, 28908, 28909, 2891, 2892, 2893, 2894, 2895, 2896, 2897, 28970, 28971, 28979, 2898, 28980, 28981, 28989, 2899, 571, 5710, 5711, 5712, 5713, 5714, 5715, 57150, 57151, 57152, 57158, 57159, 5716, 5718, 5719, 790, 7900, 7901, 7902, 7903, 7904, 7905, 7906, 7907, 7908, 7909.

**ICD10**: B20, B21, B22, B23, B24, C40, C41, C81, C82, C83, C84, C85, C86, C87, C88, C89, C90, C91, C92, C93, C94, C95, C96, D45, D46, D47, D55, D56, D57, D58, D59, D60, D61, D63, D640, D641, D642, D643, D644, D65, D66, D67, D68, D69, D70, D71, D72, D73, D74, D75, D76, D77, D80, D81, D82, D83, D84, D85, D86, D87, D88, D89, K70, K71, K74, R70, R71, R72, R73, R74, R75, R76, R77, R78, R79
